# A tool for CRISPR-Cas9 gRNA evaluation based on computational models of gene expression

**DOI:** 10.1101/2024.06.08.598047

**Authors:** Shai Cohen, Shaked Bergman, Nicolas Lynn, Tamir Tuller

## Abstract

CRISPR based technologies have revolutionized all biomedical fields as it enables efficient genomic editing. These technologies are often used to silence genes by inducing mutations that are expected to nullify their expression. To this end, dozens of computational tools have been developed to design gRNAs, CRISPR’s gene-targeting molecular guide, with high cutting efficiency and no off-target effect. However, these tools do not consider the induced mutation’s effect on the gene’s expression, which is the actual objective that should be optimized. This fact can often lead to failures in the design, as an efficient cutting of the DNA does not ensure the desired effect in protein production. Therefore, we developed EXPosition, a computational tool for gRNA design. It is the first tool designed to improve the true objective of using CRISPR: the effect it has on gene expression. To this end, we used predictive deep-learning models for the relevant gene expression steps: transcription, splicing, and translation initiation. We validated our tool by demonstrating that it can classify sites as “silencing” or “non-silencing” better than models that consider only the cutting efficiency. We believe that this tool will significantly improve both the efficiency and accuracy of genome editing endeavors. EXPosition is available at http://www.cs.tau.ac.il/~tamirtul/EXPosition.

## INTRODUCTION

Over the past decade, significant progress has been achieved in the field of genome editing, largely attributed to the utilization of CRISPR (Clustered Regularly Interspaced Short Palindromic Repeats) and its Cas (CRISPR-associated) proteins (reviewed in (1)). The Cas9 protein creates double-stranded breaks (DSBs) that are subsequently repaired by the cell’s repair mechanisms, usually through NHEJ (non-homologous end joining), leading to the potential introduction of indels. One of the areas where these advancements have occurred pertains to gene silencing, with the primary objective being the selective inhibition of a specific gene without affecting others. Various methods for gene silencing using CRISPR exist, including expression inhibition through CRISPRi, the introduction of point mutations, and the insertion of premature stop codons. Another common approach involves the use of Cas9 proteins to induce mutations in start codons, thereby inhibiting translation initiation (2–4).

In most computational models predicting CRISPR activity, researchers have shown interest in the DSB’s location and likelihood, as well as the identity of the resulting mutation (5–23). This paradigm assumes that if a mutation is induced, then the gene’s expression will significantly decrease; however, this is not always the case. For example, a gene with a mutated start codon can still be translated due to an alternative in-frame start codon, so the resulting protein remains functional (24) (Fig. 1A).

**Fig 1:**
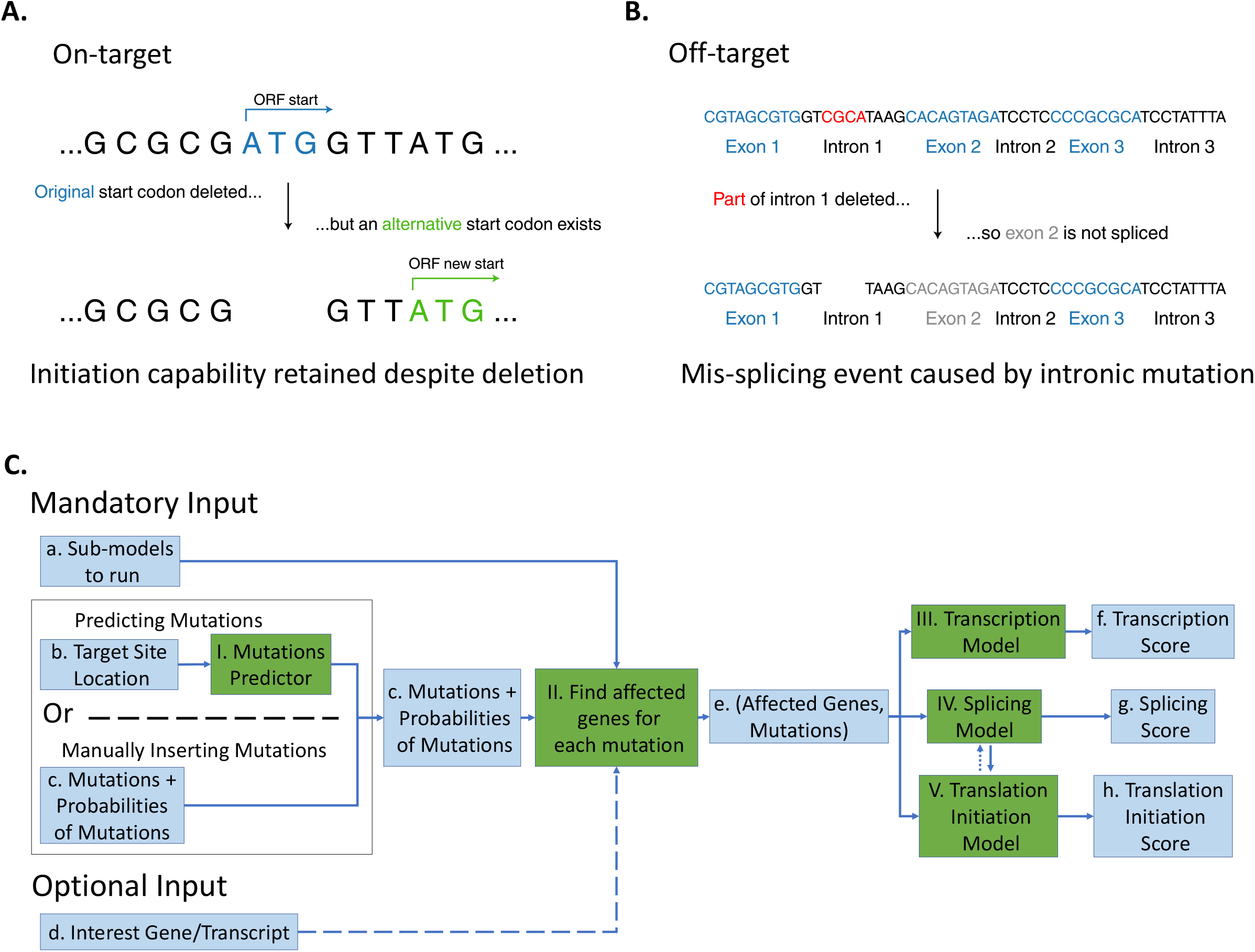
Overview of the tool’s rationale and structure. **A**. A given gene’s canonical start codon (blue) is deleted by a CRISPR mutation, to silence the gene. However, since an alternative ATG in the same reading frame exists (green), translation initiation still occurs, effectively keeping the gene unsilenced. **B**. Part of intron 1 (red) is deleted by a CRISPR mutation; although each exon’s pre-mRNA sequence remains intact, the deletion causes a missplicing event, resulting in exon 2 not being spliced; Thus, the mRNA includes only exons 1 and **C**. The EXPosition pipeline. First, the tool either predicts the most likely mutations induced in a CRISPR target site or receives specific mutations from the user; it then searches for potentially affected genes and assesses the mutation’s effect on their splicing, transcription, and translation initiation. The user can also insert specific genes/transcripts for analysis, thus disregarding other genes/transcripts which are affected by the mutations. Solid/dashed arrows represent mandatory/optional input, respectively.

Furthermore, transcription and splicing are influenced by various properties of the DNA sequence (25, 26), and not every change to the original sequence will result in a measurable change in the gene’s expression. Therefore, mutated off-target genes, i.e., genes mutated by CRISPR even though the gRNA wasn’t meant to target them, do not necessarily have their expression affected. On the other hand, even if each exon’s DNA sequence remains unmutated, the gene’s expression could be affected by intronic or intergenic mutations (Fig. 1B). Thus, mutations in all positions require careful evaluation to determine whether they exert a discernible effect on expression.

Here we present a tool that addresses these issues by explicitly considering CRISPR’s effect on the target site’s phenotype rather than the genotypic change. Our tool predicts the impact of CRISPR’s action on three aspects of gene expression: transcription, splicing, and translation initiation. Since not every mutation will significantly affect gene expression, researchers using our tool can save time and money when deciding which gRNA is most likely to achieve the desired change in gene expression (e.g. silencing) without affecting the expression of off-target genes.

## MATERIAL AND METHODS

### General pipeline of the tool

Our tool, called EXPosition, accepts a CRISPR target site location and predicts the phenotypical effect of CRISPR at that site, i.e. the effect of the induced mutation on gene expression. The tool’s modules are summarized in Fig. 1C. Firstly, the user chooses which sub-models to run (Fig. 1C:a): transcription, splicing and translation initiation. Then, the user inputs a target site location (Fig. 1C:b). Using CRISPRedict (27) and Lindel (28), the tool predicts the most likely mutations to be induced by CRISPR, along with their probabilities (Fig. 1C:I). Alternatively, the user can specify the mutations and their probabilities (Fig. 1C:c). In both cases, these mutations are then analyzed in the chosen sub-models. In each sub-model (Fig. 1C:III-V), we evaluate the effect of each mutation on one aspect of expression for each human gene, thereby finding the genes affected by the mutation. We then multiply each mutation’s phenotypic score by the probability of the mutation, and sum over all mutations to arrive at an expected value of effect for that aspect of expression for a certain gene (Fig. 1C:f-h). If multiple genes have been affected by the mutations, the gene that received the highest score (i.e., most detrimental effect on expression) will be the output of the sub-model. Thus, the final score for each sub-model would be:

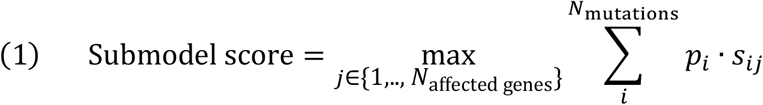

Where *j* ranges from 1 to *N*_affected genes_, which is the number of genes affected by the predicted mutations and is usually 1-2 (Fig. 1C:e); *N*_mutations_ is the number of top mutations (i.e. most probable mutations) considered (by default, *N*_mutations_ = 4); *p*_*i*_ is the probability of mutation *i* occurring; and *s*_*ij*_ represents the expected effect of mutation *i* on gene *j* according to the sub-model, i.e., its effect on transcription, splicing or translation initiation. *s*_*ij*_ is normalized to range between 0 and 1 in the following way:

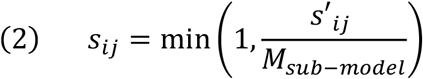

Where *s′*_*ij*_ represents the highest raw sub-model score for mutation *i* over all gene *j*’s transcripts, and *M*_*sub−model*_ is the maximal score predicted by the sub-model on all ClinVar mutations (see section “High-scoring mutations are overrepresented in ClinVar-designated pathogenic mutations”).

Additionally, instead of evaluating every human coding gene/transcript affected by the mutations, the user can input a gene or transcript of interest (Fig. 1C:d); if they do so, the mutation’s effects in the selected sub-models will be checked only against that gene or transcript. If a transcript of interest was set, *s′*_*ij*_ is the score of that transcript (instead of the maximal score over all the associated gene’s transcripts).

In the transcription sub-model (Fig. 1C:III), the score (*s*_*ij*_) represents the predicted relative change in mRNA levels caused by a mutation; this change is predicted using Xpresso (25). In the splicing sub-model (Fig. 1C:IV), the score signifies the predicted loss of functionality of the gene’s proteins (based on evolutionary conservation) caused by a mutation, while considering any mis-splicing events and changes in the position of their start codon; this is accomplished using Oncosplice (29). In the translation initiation sub-model (Fig. 1C:V), the score denotes the predicted relative change in the start codon’s efficiency of translation initiation, while considering any mis-splicing events; this is achieved using TITER (30). The following sections detail the different parts of EXPosition.

### Predicting the genotypic outcome of CRISPR’s DSB

As a first step, based on the user’s cut site location, we extract the target site along with its flanking sequences to predict the DSB’s probability and resulting indels using CRISPRedict (27) and Lindel (28) respectively (Fig. 1C:I). CRISPRedict is a linear regression model that predicts the probability of cutting by sgRNAs; it takes as input a 30nt-long sequence surrounding the cut site: 4nt upstream to the cut site, 20nt of the site, and the following 6nt downstream to site. Lindel is a logistic regression model that predicts the likelihood of NHEJ mutations induced by CRISPR; its input is a 60nt sequence centered around the cut site, and its output consists of the predicted probabilities of 557 possible mutations: deletions around the cut site of up to 30nt, every possible insertion of 1-2nt, as well as a single collective mutation for any insertions of *≥*3nt.

We analyze only the *N*_mutations_ (by default *N*_mutations_ = 4) most likely mutations predicted by Lindel in our tool, as checking all possibilities isn’t feasible timewise. We normalize the probabilities of these mutations so that their sum equals 1. We also exclude insertions longer than 2nt, as Lindel does not provide explicit mutations for such cases, which have been demonstrated to be exceedingly rare (23, 28).

Thus, the probability for each mutation is calculated as follows:

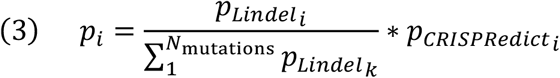

Where 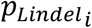 and 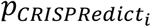 represent the probabilities from Lindel and CRISPRedict of mutation *i*, respectively; and *N*_mutations_ is the number of most probable mutations taken from Lindel.

The predicted mutations and their probabilities serve as input for the three sub-models described in the following sections. Alternatively, users can manually input specific mutations of interest along with their probabilities, which can include both indels and substitutions.

### Transcription sub-model

To predict the effect of a mutation on transcription (Fig. 1C:III) we employ Xpresso (25), a deep learning model (Fig. S1) that predicts mRNA steady-state abundance based on the nucleotide context around a Transcription Start Site (TSS). More details can be found in supplementary section 1. Its input consists of a 10.5kb context around the TSS, while the output is the log_10_ mRNA expression level of the respective gene (Fig. 2A).

**Fig 2:**
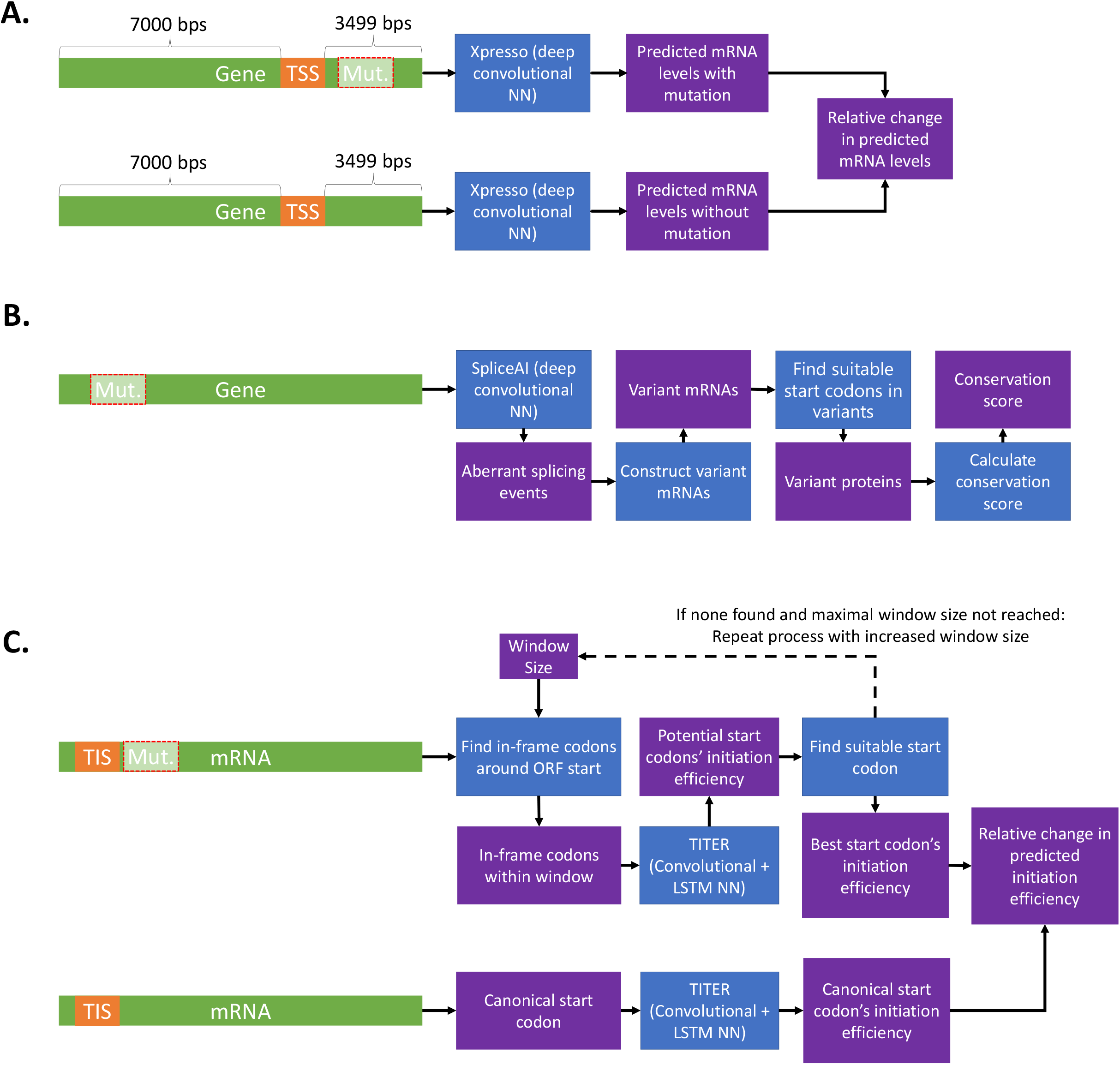
Overview of the gene expression sub-models. **A**. The transcription model takes as its inputs the sequences surrounding the Transcription Start Site (TSS), which include both the mutated and unmutated sequences. Specifically, it considers 7000 base pairs upstream and 3499 base pairs downstream of the TSS. The mutated and unmutated sequences are each fed into Xpresso, which then predicts the mRNA levels for the mutated and unmutated genes. The model’s output is the relative change in mRNA levels compared to the unmutated gene. **B**. The splicing model receives the gene’s predicted mutations and uses them in SpliceAI to identify potential aberrant splicing events. Accounting for any mis-splicing events detected, variant mRNAs are generated. These variant mRNAs are subsequently assessed by our Translation Initiation model to identify a suitable start codon. The variant mRNA sequences are translated into amino acids and employed to calculate the gene’s conservation score. **C**. The Translation Initiation model accepts both the mutated and unmutated transcripts as inputs. In an iterative procedure, the mutated transcript is searched for in-frame codons within a defined window. These codons’ initiation capabilities are predicted by TITER. If a suitable codon is identified, defined as one with better efficiency than the canonical start codon or ranking within 5% of its efficiency when compared to all human transcripts, it is selected and returned. If no suitable start codon is found in the initial window, the iterative process continues with an expanded window size. This process continues until a suitable start codon is discovered or until a maximum window size is reached, at which point the best available codon is returned. Simultaneously, the unmutated mRNA’s start codon undergoes analysis by TITER, and its efficiency ranking is used to calculate the relative change in initiation efficiency ranking between the canonical start codon and the best new start codon identified in the mutated sequence.

For a given mutation *i* and gene *j*, we examine whether the mutation could potentially impact the gene’s transcription levels by considering its transcripts’ TSSs and checking if the mutation falls within 7kb upstream or 3.5kb downstream of them (i.e. in the region that Xpresso considers when evaluating a TSS). For each potentially impacted transcript, we calculate the Xpresso score of the 10.5kb sequence around its TSS before and after the mutation (denoted *r*_*WT*_ and *r*_*mutated*_, respectively). Each transcript’s final transcription score reflects the relative change in mRNA transcription levels following the mutation:

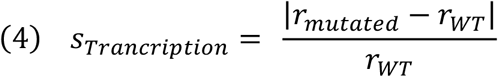

*s′*_*ij*_ in Eqn. 2 is the maximal *s*_*Trancription*_ caused by mutation *i* over gene *j*’s transcripts. If a specific transcript of interest was provided, the output score pertains solely to that transcript.

### Finding mRNA isoforms following a mutation

Both the splicing model (Fig. 1C:IV) and the translation initiation model (Fig. 1C:V) analyze isoforms of transcripts following mutations and potential aberrant splicing events. We obtained splice site annotations by Ensembl (31). To predict mis-splicing events, we utilize SpliceAI (26), a deep learning tool (Fig. S2) that predicts the change in a position’s probability to function as a splicing donor/acceptor site following a mutation. More information can be found in supplementary section 2. We use these annotated splice sites and the splicing changes predicted by SpliceAI to generate all possible mRNA isoforms following the mutation by concatenating donor and acceptor splice sites (Fig. S3). Further details are available in supplementary section 3.

### Splicing sub-model

The splicing sub-model (Fig. 1C:IV) assesses the impact of a mutation on a gene’s viability by examining the isoforms generated for each of the gene’s transcripts. To gauge the effect of a mutation on a protein’s functionality we use Oncosplice (29). This model receives a mutation and a gene as input and predicts how much the mutation disrupts the gene’s protein function. This disruption is scored using evolutionary conservation information (Fig. 2B).

For each isoform, a sliding window is employed to identify the most conserved area that is affected by the mutation; the window’s length is set to the average domain length of all human proteins. The score of each transcript is the average score of its isoforms (Fig. S4); and finally, the gene’s score is the maximal transcript score, i.e. the transcript whose function was most significantly disrupted. If a specific transcript of interest was provided, the output score pertains solely to that transcript. For further information, please refer to supplementary section 4.

### Translation initiation sub-model

The translation initiation sub-model (Fig. 1C:V, Fig. 2C) assesses the ability of mutant variants to initiate translation by searching for suitable start codons within the isoforms identified by the splicing model (see section “Finding mRNA isoforms following a mutation”). The translation initiation score of the suitable codons is determined using TITER (30), a deep learning tool (Fig. S5) that integrates a deep learning algorithm with known codon compositions of Translation Initiation Sites (TISs) to predict TIS functionality. For additional information, please consult supplementary section 5.

For each isoform, we locate the start of the coding sequence through local alignment with the original transcript’s coding sequence’s start. An iterative process is then initiated where TITER examines all in-frame NUG and ACG codons (where N can be any nucleotide) within a window surrounding the coding sequence’s start, with the window size increasing in each iteration. The process concludes when either the best new codon is discovered (with a TITER score sufficiently close to that of the canonical start codon), or when the maximum window size is reached. Further information can be found in supplementary section 6. We then calculate the WT start codon’s TITER score rank, compared to the TITER scores of all human canonical start codons; this rank is denoted as *i*_*WT*_. We repeat this calculation for the best new start codon found for the isoform, whose rank is denoted *i*_*mutated*_. The isoform’s score is defined as the relative change in the isoform start codon’s TITER score rank:

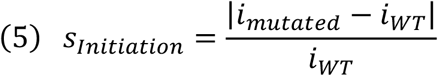

Like the splicing sub-model, we calculate the transcript’s initiation score as the average of the isoform scores; and the gene’s initiation score (i.e. s*′*_*ij*_ from Eq. 2) as the highest score among all its transcripts (Fig. S4). Likewise, if a specific transcript of interest is provided, we provide the score exclusively for that transcript.

### High-scoring mutations are overrepresented in ClinVar-designated pathogenic mutations

We want to inform the user, for each sub-model, how detrimental a score is to the model’s corresponding aspect of a gene’s expression. Thus, we used our tool to analyze mutations from the ClinVar dataset (32), which contains mutations and their phenotypes accumulated from laboratories and researchers globally. We analyzed ~325k mutations tagged as benign (192k) or pathogenic (133k).

For each EXPosition sub-model, we examined the 3254 mutations in each percentile range (i.e. 99%-100%, 98-99% etc.) and calculated the fraction of pathogenic mutations in that set (Fig. 3; the score thresholds for each percentile are detailed in supplementary section 7, Table S1). We denote this fraction *S*_*ClinVar*_. A higher fraction indicates better recognition of pathogenic mutations by the sub-model. All sub-models provided meaningful rankings on the ClinVar dataset from a certain threshold. The thresholds of the transcription/splicing/translation initiation sub-models correspond to the 92/67/97 percentiles. *S*_*ClinVar*_ values for the higher percentiles exhibited significant enrichment of pathogenic mutations in almost all top 5 percentiles (*p* < 10^*−*25^, *p* < 10^*−*324^, *p* < 10^*−*308^ for the transcription, splicing and translation initiation sub-models respectively using the hypergeometric test).

**Fig 3:**
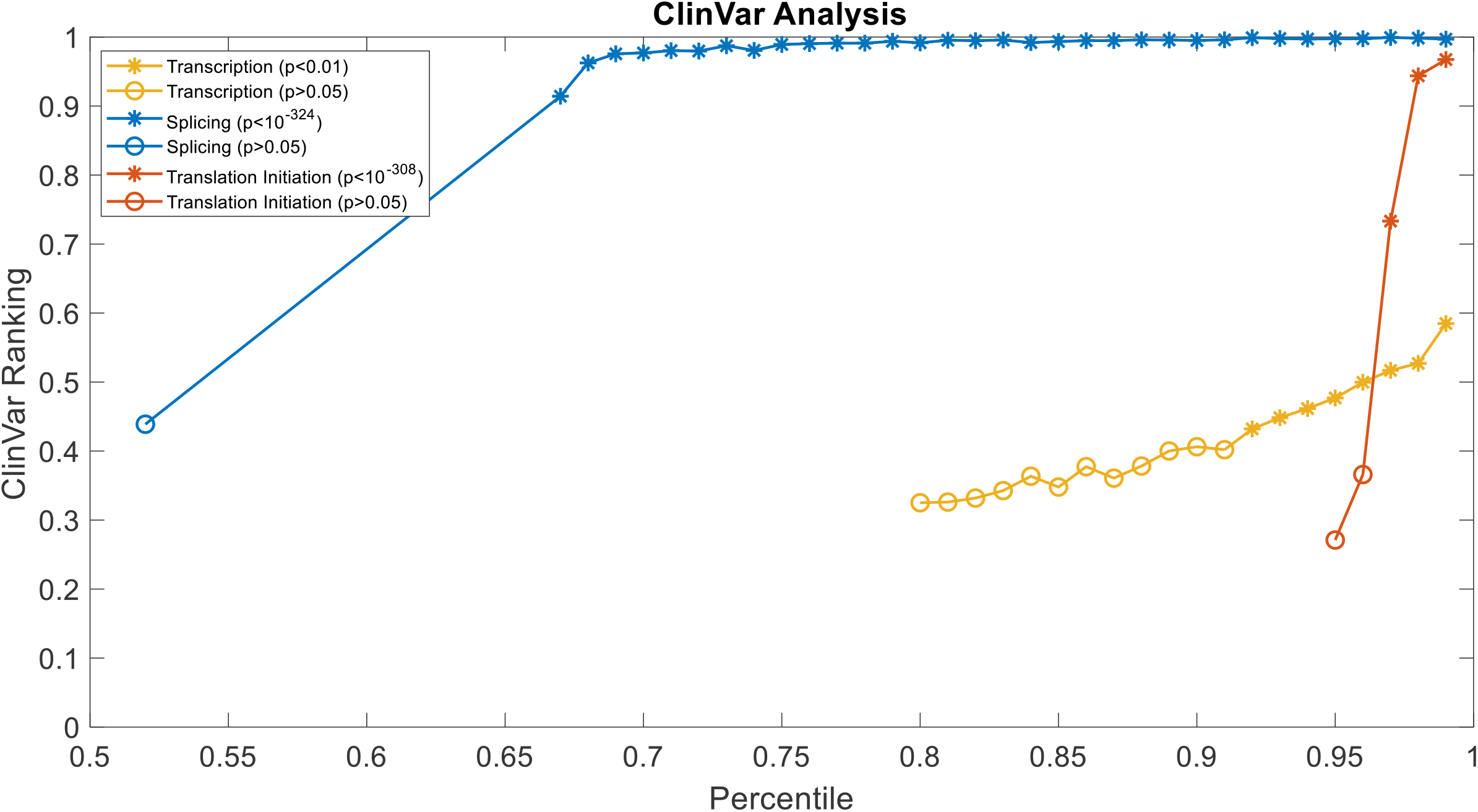
ClinVar analysis. S_ClinVar_ values of the transcription, splicing, and translation initiation sub-models (yellow, blue, orange lines and dots, respectively) for different percentiles. Asterisks denote significant p-values (p < 0.05) and empty circles denote non-significant p-values (p ≥ 0.05). The highest p-value (i.e. closest to 0.05) calculated for each sub-model is listed in the legend. All p-values were corrected using FDR. Note that for each sub-model, the dots begin at the first percentile which corresponds to a non-zero score, meaning that 52/80/95% of mutations received a score of 0 in the transcription/splicing/translation initiation sub-models respectively.

Following this analysis, each sub-model’s ClinVar threshold was set as the lowest percentile in which we observed a significant enrichment of pathogenic mutations (e.g. the 97^th^ percentile for the translation initiation sub-model). We then calculated the maximal score predicted by each sub-model for the whole set of ClinVar mutations and normalized the thresholds by these maximal values. To assess the impact of a gRNA on a specific expression aspect, we compute an average score across the corresponding sub-model for all predicted mutations, weighted by their probabilities. If the mean sub-model score surpasses its designated threshold, EXPosition informs the user that this aspect is affected. Alternatively, if a mutation was inputted manually, its score is checked against the sub-model’s threshold and the user is notified accordingly. The maximal scores from the ClinVar analysis for each sub-model used to normalize the scores (Eq. 2), as well as the raw and normalized final thresholds, can be found in supplementary section 7, Table S2.

### Analysis of data from HT29 cells

We conducted an analysis on data comprising sgRNAs and their impact on cell fitness, sourced from Doench et al. (33). In this analysis, we employed SVM classifiers to differentiate between sites that were most likely to silence their target gene and those least likely to do so. You can find further details in the section titled “EXPosition Improves the Prediction of the Effect of CRISPR Editing on Cell Fitness Compared to Solely Considering Cutting Efficiency” in the Results.

The analysis carried out on the HT29 cell data did not reveal any statistically significant difference between a classifier that solely considered cutting efficiency and one that incorporated both efficiency and EXPosition. We attribute this lack of distinction to the heightened noise present in the HT29 data, as compared to the noise observed in the A375 data (as illustrated in Fig. S6). This noise became evident when attempting to predict measurements from one of the experiment’s repeats to another, as detailed in supplementary section 9

## RESULTS

### EXPosition is a tool to predict a mutation’s effects on gene expression

We have developed EXPosition (EXPosing CRISPR’s Impact on EXPression and Position), a computational tool designed to assess CRISPR sites based on their impact on three key stages of gene expression: transcription, splicing, and translation initiation. The tool begins by predicting the most probable mutations and their associated probabilities resulting from CRISPR utilization. Subsequently, EXPosition assesses the impact of each mutation on these three aspects of gene expression and outputs a score for each gene affected by the mutations.

Our tool (depicted in Fig. 1C) employs deep-learning algorithms, all of which have undergone independent validation, to execute these tasks. The transcription sub-model (Fig. 1C:III) predicts the relative change in mRNA levels following the mutation. As for the splicing and translation initiation sub-models (Fig. 1C:IV-V), we consider all possible isoforms generated by the mutation through alternative splicing. These isoforms can lead to a distinct protein from the original one due to the following factors:

1. **Aberrant splicing events:** These can arise by either creating new donor/acceptor sites or deleting existing ones, resulting in an altered splicing pattern of the mRNA.
2. **Alterations in initiation site usage:** Mutations in the start codon or its context can modify the initiation capability. Factors such as the nucleotide context and folding energy of the start codon play pivotal roles in determining initiation efficiency (as reviewed in (34)). Any change in these aspects could impact the initiation capability of the original start codon. Additionally, other potential start codons, such as ATGs in the same reading frame, could serve as alternative initiation sites, preserving the transcript’s initiation capability (35) and potentially retaining the protein’s function. Consequently, an assessment of the initiation capability of all potential start codons, including the original ATG if unaltered, is necessary to predict whether translation initiation can occur.
3. **Elimination of the gene’s stop codon:** This type of mutation can lead to the addition of potentially unnecessary amino acids to the translated protein.

Therefore, our tool comprehensively evaluates the potential effects of mutations on these three aspects of gene expression, utilizing deep-learning algorithms that have undergone rigorous validation.

The mRNA’s isoforms are constructed by predicting the mutation’s effect on alternative splicing and concatenating relevant exons, i.e. exons with viable splicing donor/acceptor pairs. These isoforms are then passed to the Splicing and Translation Initiation sub-models, which assess the viability of these isoforms based on amino acid conservation and the translation initiation capability of the isoforms, respectively. Details regarding each sub-model appear in the “Material and methods” section. The tool is written in Python 3.9 and can be accessed in http://www.cs.tau.ac.il/~tamirtul/EXPosition. The tool’s GUI is shown in Fig. 4.

**Fig 4:**
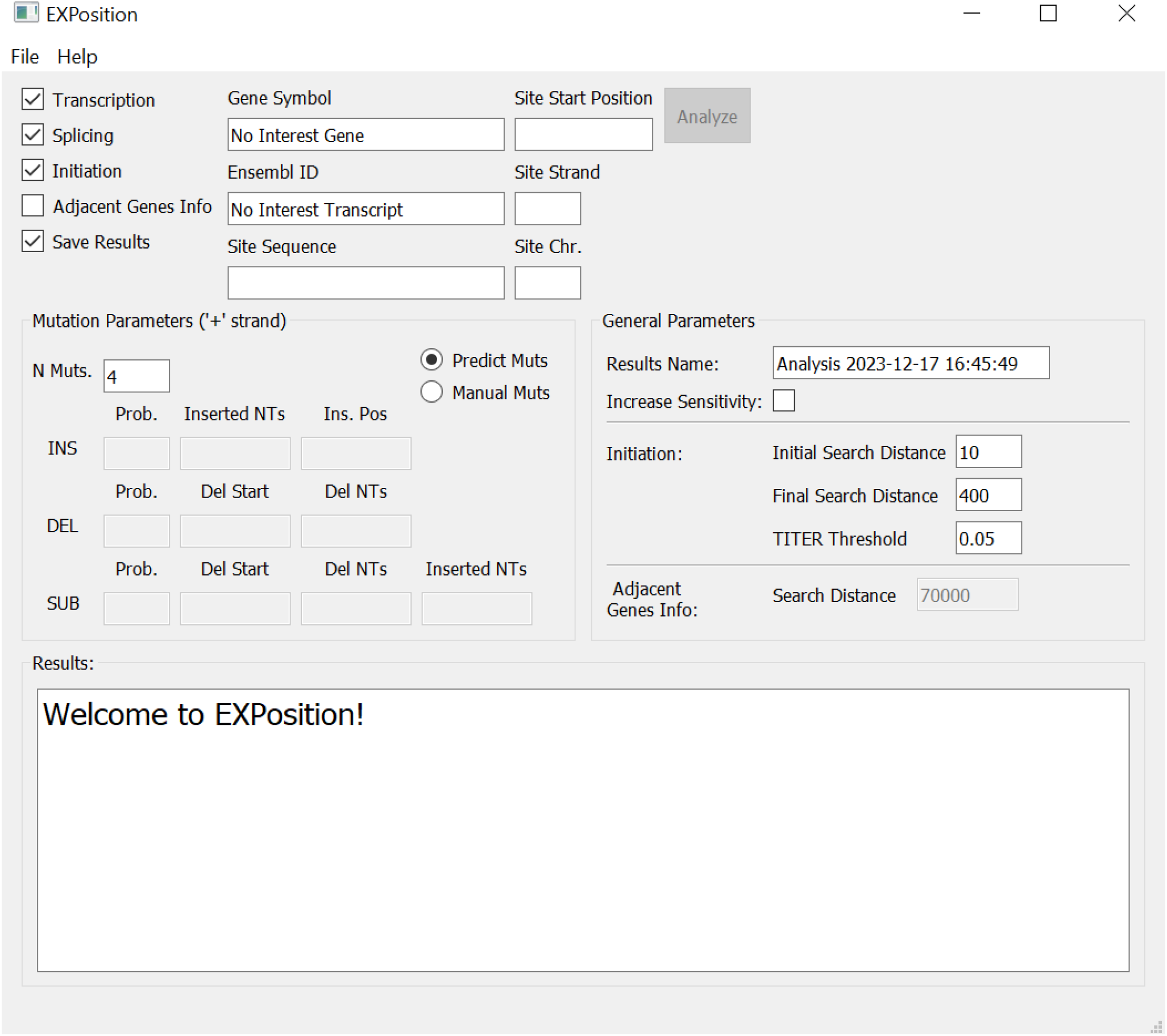
The GUI of EXPosition. The user needs to check the boxes of the sub-models to be run (Transcription/Splicing/Initiation, upper left corner) and provide the target site information (Chromosome, strand, position andff 20nt sequence). If the user predicts the resulting mutations via our tool, “Predict Muts” should be selected; otherwise, the user can insert the mutations manually and select “Manual Muts”. The initiation sub-model parameters can be adjusted manually by the user. The results are saved in a csv file and shown in the Results textbox.

### EXPosition improves the prediction of the effect of CRISPR editing on cell fitness in comparison to only considering cutting efficiency

To validate our tool, we searched for datasets containing measurements of gene expression following CRISPR editing. We selected the data published by Doench et al. (33); In their study, the authors performed a negative selection screening using sgRNAs that targeted multiple sites with the aim of identifying essential genes in A375 and HT29 cell lines.

They infected the cells using lentiviruses, causing them to express sgRNAs and Cas9, and measured the fold change in sgRNA levels following CRISPR’s action (Fig. 5A).

**Fig 5:**
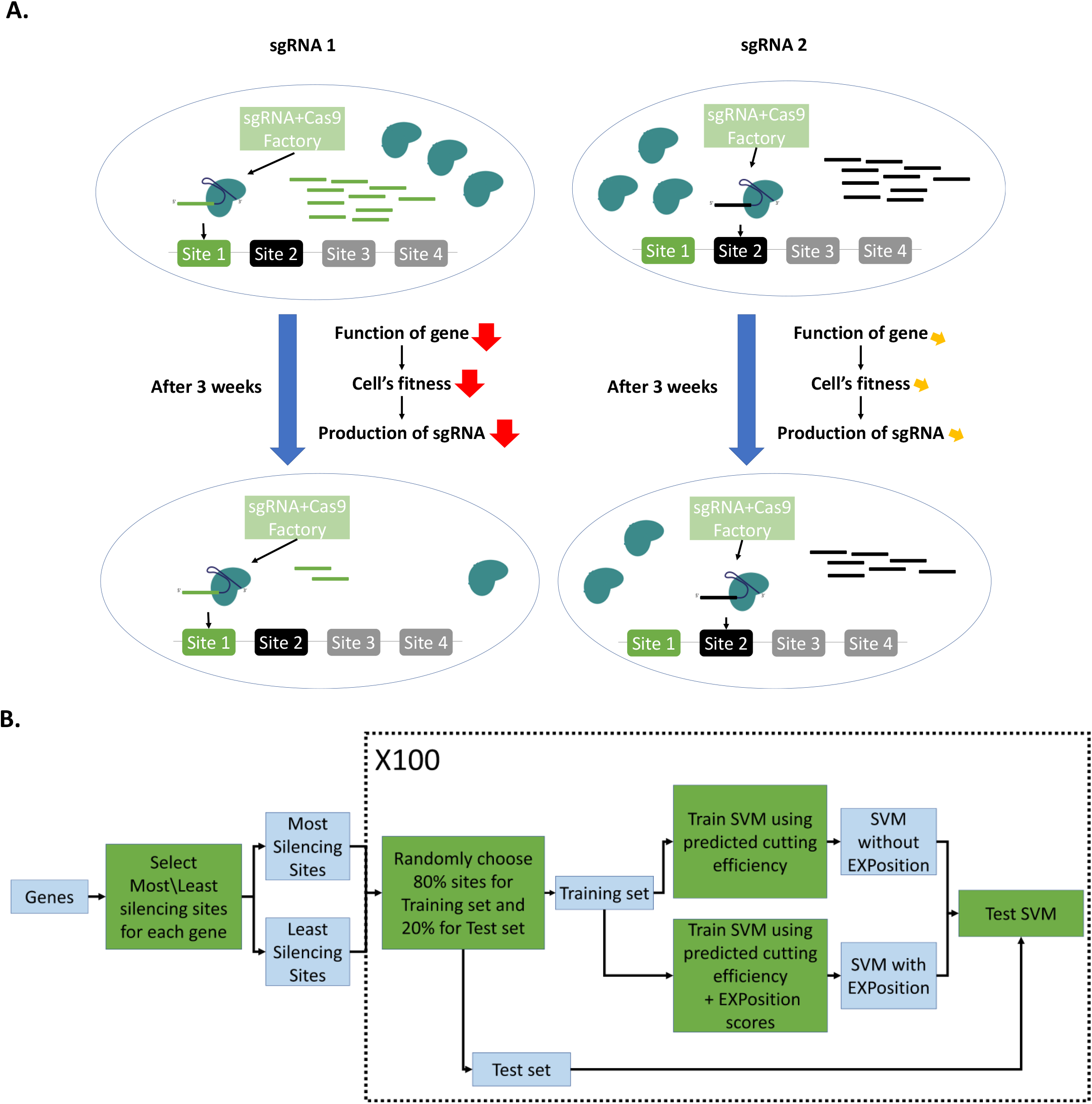
Illustration of the Doench al. experiment and analysis. **A**. The cells in the top row are infected with a lentivirus and produce Cas9 proteins and sgRNAs; the left/right cell produces sgRNA1/2, which affects site 1/2 in a given gene, respectively. After 3 weeks (bottom left), sgRNA1 induces mutations in the gene, which affect its function and cause the cell’s fitness to decrease; this results in lower amounts of sgRNA1 being produced. Meanwhile, sgRNA2 induces mutations in the gene as well, but its function – and the cell’s fitness – is minimally affected; thus, sgRNA2 levels stay nearly the same. Each gene in the dataset contains 4-6 different target sites. **B**. We conducted 100 iterations of Monte-Carlo cross validation, training two SVM classifiers with/without EXPosition, each time taking a random 80% of the data and testing the models on the remaining 20%. See full details in the main text.

Their underlying assumption was that sgRNAs targeting essential genes would negatively impact the cell’s fitness, leading to a reduction in the production of these sgRNAs. We believe that this assumption generally holds true to some extent for any gene (36, 37). Furthermore, we believe that this relationship is, to some extent, attributable to alterations in gene expression. Consequently, we anticipated observing a stronger correlation between the impact on fitness and the predicted effect on gene expression than between the impact on fitness and the efficiency of DNA cutting alone.

The Doench dataset (33) encompassed genes with 4-6 different target sites, each targeted by a distinct sgRNA. Our hypothesis posited that, for a given gene, the sgRNA that had the most pronounced effect on the gene’s expression would also exhibit a significantly more negative effect on the cell’s fitness when compared to the sgRNA with the least effect on the gene’s expression.

The dataset comprised two repetitions of the experiments conducted using two distinct delivery systems (lentiGuide and lentiCRISPRv2) in the A375 cells, as well as three repetitions for a single delivery system in the HT29 cells. In both the A375 and HT29 cells, we calculated the average values across these repetitions, as outlined in further detail in supplementary section 8.

For each of the 18,086 genes within the dataset, we selected two sites: the most silencing site, denoted as the site with the largest reduction in sgRNA levels; and the least silencing site, characterized by the smallest decrease in sgRNA levels. We labeled these sites as “silencing” and “non-silencing,” respectively. Subsequently, we applied our model to each of these sites, predicting CRISPR’s impact on transcription, splicing, and translation initiation.

To evaluate whether EXPosition can effectively distinguish between silencing and non-silencing sites, we trained two SVM classifiers designed to categorize a given site as either silencing or non-silencing. The first classifier was trained exclusively using the predicted cutting efficiency (utilizing CRISPRedict) as a feature, while the second classifier incorporated both the predicted cutting efficiency and the 3 EXPosition sub-model scores as features (as illustrated in Fig. 5B). We conducted training for both classifiers on randomly selected 80% of the sites (29k sites) and subsequently tested the trained classifiers on the remaining 20% of the sites (7k sites). This process was repeated 100 times, with the training and test sets randomly selected for each iteration.

We then calculated each SVM model’s loss on its test set, thereby acquiring two loss distributions: loss from models using only cutting efficiency, and loss from models using efficiency and EXPosition. The losses achieved by models using efficiency and EXPosition were significantly lower than models using only efficiency (p = 2×10^−18^, Wilcoxon rank sum test; Fig. 6A). As a control, we repeated the Monte Carlo cross-validation process, this time shuffling the site labels (silencing and non-silencing) before training. In this case we found that the loss distributions from both classifiers were similar and overlapping, with no statistically significant difference (p = 0.17, Wilcoxon rank sum test; Fig. 6B).

**Fig 6:**
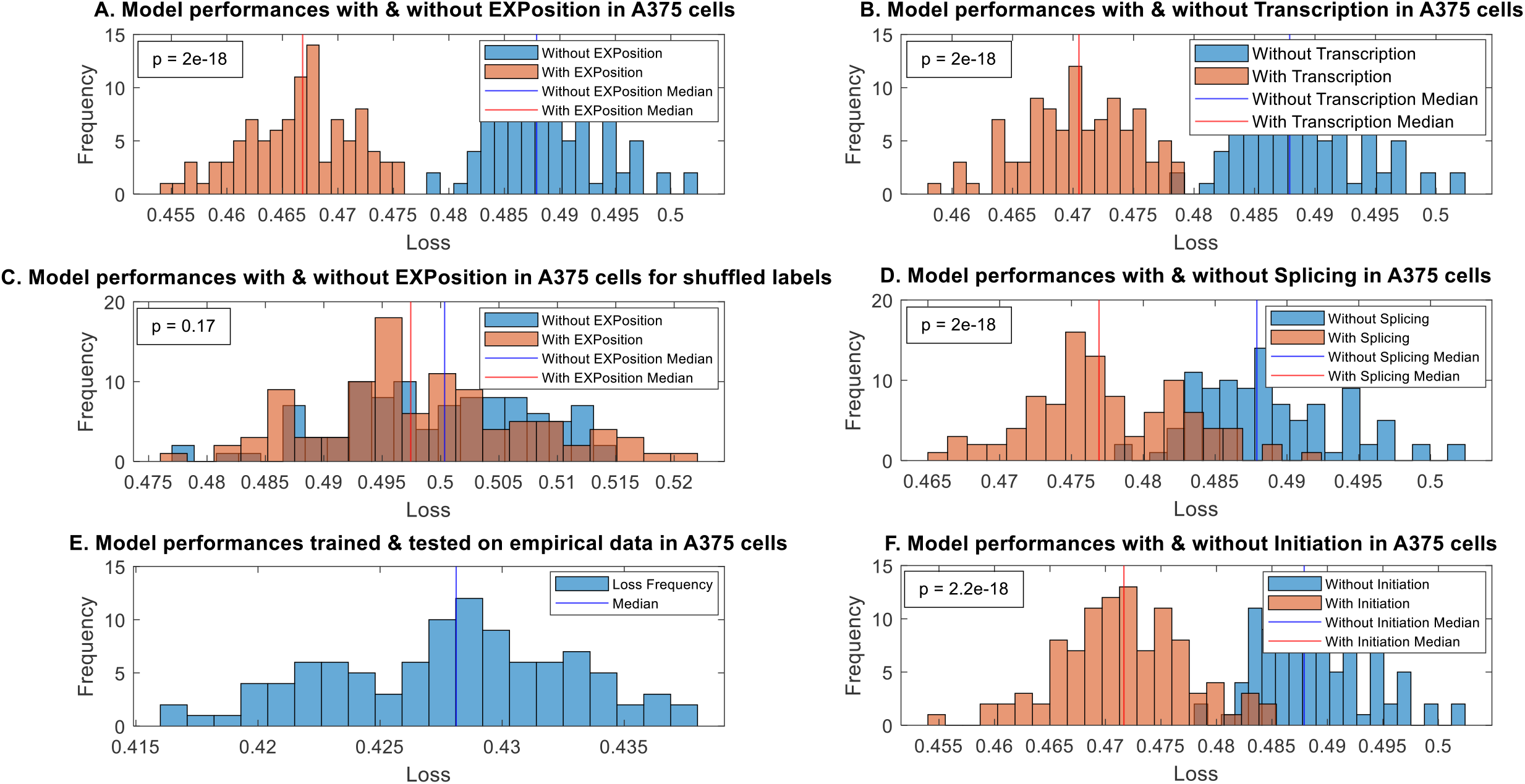
Using EXPosition with cutting efficiency significantly improves classification of silencing/non-silencing sites. Histograms of SVM classifier loss, from models trained using only cutting efficiency (blue) and models using cutting efficiency and the outputs from EXPosition’s models (red). Vertical lines represent distribution median. p-value calculated using Wilcoxon rank sum test **A-B**. Histograms of SVM classifiers trained with/without the 3 EXPosition scores, using the A375 original data (A) or a dataset where the site labels were shuffled in training (B). **C**. Results from 100 SVM classifiers trained using results from one infection method (lentiGuide), where the most/least silenced sites for each gene were chosen using the results from the other infection method (lentiCRISPRv2). **D-F**. Histograms of SVM classifiers trained with/without the transcription (D), splicing (E) or translation initiation (F) sub-model score, using the A375 original data.

It is important to emphasize that the predictive power of our model (loss of 0.47; Fig. 6B) is similar to the predictive power of using one repeat of the experiment to predict the second repeat of the experiment (loss 0.43; Fig. 6C). Thus, it is very significant and likely to be close to what we can get for these experimental data (see more details supplementary section 9).

We also validated that each one of our features contributed to the prediction, by repeating the process described in Fig. 5A using only one sub-model of EXPosition at a time, i.e. comparing models trained on efficiency to models trained on efficiency and either transcription, splicing or translation initiation (Fig. 6D-F); the difference between the loss distributions was significant in all 3 cases, demonstrating that each of the models contributed to the classification process, and is therefore valid in and of itself.

### Examples of silencing sites found with EXPosition

Finally, we provide a few examples where we believe our tool can better predict which site is more likely to silence a gene, compared to just using cutting efficiency (Table 1, Fig. 7). We use the Doench dataset and compare the most silenced target site of a gene to the least silenced site, as defined in section “EXPosition successfully differentiates between the least and most silencing sites” and Fig. 4A. In each example, examining only the cutting efficiencies would point to the wrong site as most silencing.

**Table 1:** Examples of sites successfully differentiated into least-silencing and most-silencing. In each case the predicted cutting efficiency of the least silenced site is higher than the most silenced site’s efficiency. The cells marked in green are where our tool predicted a more detrimental effect on a certain aspect of the gene’s expression for the most silenced site compared to the least silenced site.

| # | Type | Gene | Measured Effect | Predicted cutting efficiency | Transcription score | Splicing score | Translation initiation score |
| --- | --- | --- | --- | --- | --- | --- | --- |
| 1 | All | SCAMP5 | Least Silenced | 0.49 | 0.018 | $9 \times 10^{-5}$ | 0 |
| | | | Most Silenced | 0.38 | 0.021 | $4.4 \times 10^{-4}$ | 0.002 |
| 2 | Transcription | MRGPRG | Least Silenced | 0.44 | 0.018 | $1.1 \times 10^{-4}$ | 0 |
| | | | Most Silenced | 0.36 | 0.034 | $6 \times 10^{-5}$ | 0 |
| 3 | Splicing | POLR1E | Least Silenced | 0.76 | 0 | 0 | 0 |
| | | | Most Silenced | 0.69 | 0 | $7 \times 10^{-4}$ | 0 |
| 4 | Translation Initiation | CCND3 | Least Silenced | 0.57 | 0 | $1.8 \times 10^{-4}$ | 0 |
| | | | Most Silenced | 0.4 | 0 | $1.8 \times 10^{-4}$ | 0.003 |

**Fig 7:**
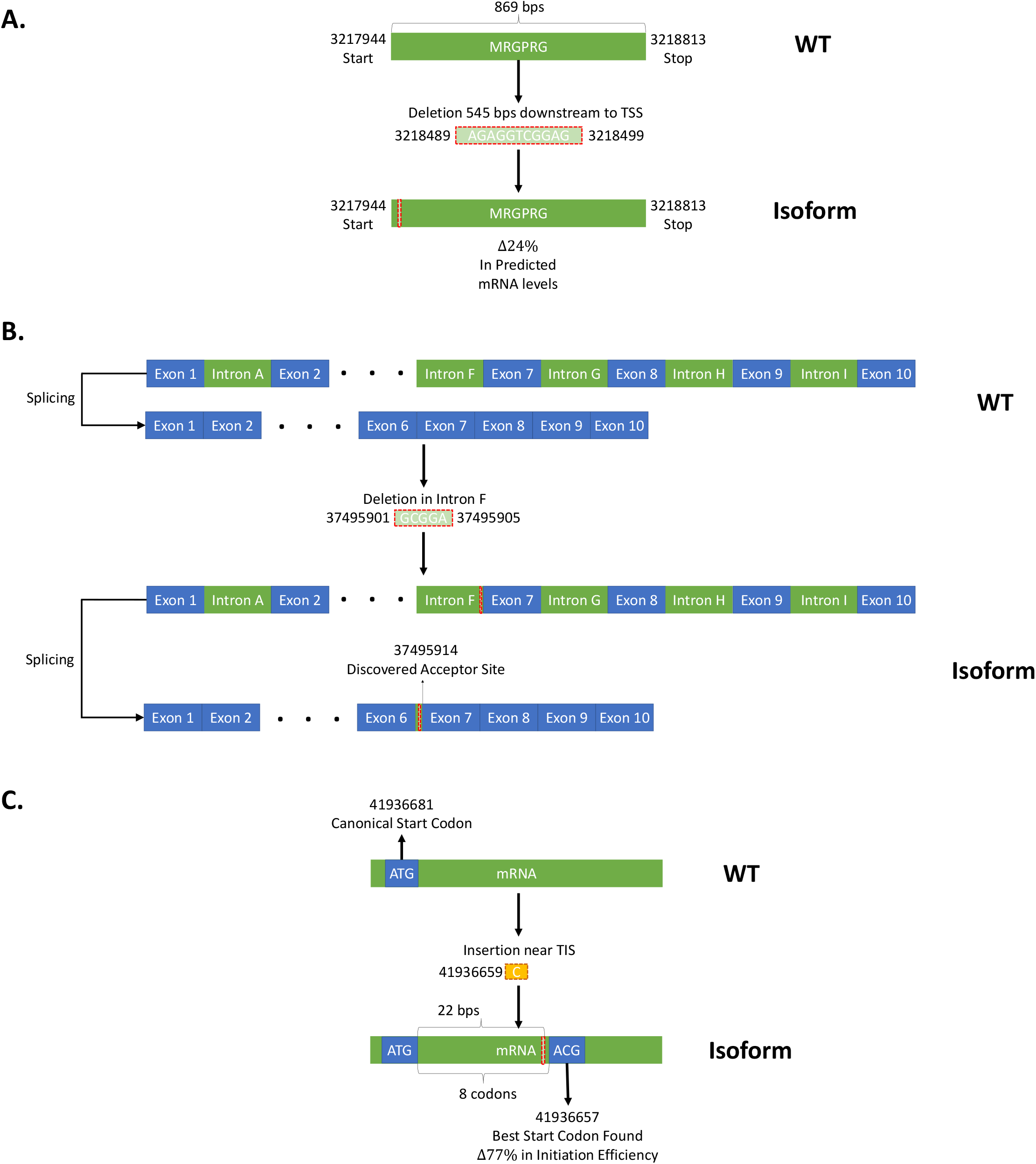
Predicted changes in gene expression following CRISPR action. Illustrations of the changes in gene expression, as predicted by EXPosition, for examples 2-4 in Table 1. **A**. One of the predicted deletions in the gene MRGPRG (which has a single coding transcript) was predicted to cause a 24% change in mRNA levels compared to the WT. **B**. A mis-splicing event is predicted to occur following a predicted deletion in the ENST00000377792.3 transcript of the POLR1E gene. A missed acceptor site causes an intron to be included in the mRNA. **C**. A predicted deletion causes a frameshift in the ENST00000415497.6 transcript of the CCND3 gene, resulting in a loss of translation initiation capability. The best alternative start codon found had a 77% change initiation ranking, relative to the ranking of the canonical start codon.

In example 1, all our sub-models grade the most silenced site with higher scores, indicating a more detrimental effect, than the least silenced site. In examples 2-4, the transcription, splicing and initiation sub-models respectively managed to correctly identify the more silencing site by themselves. In example 3, 2 out of the 4 mutations predicted by Lindel caused aberrant splicing events; in example 4, all 4 mutations were predicted to affect the translation initiation efficiency. We note that even where a sub-model did not indicate the most silenced site with a significantly higher score, the scores of the most/least silenced sites were very similar – meaning the sub-model did not heavily indicate the “wrong” site, i.e. the least silenced site, as more disruptive. Thus, each of the sub-models’ outputs, as well as their combined output, can be useful in differentiating between silencing and non-silencing sites – and in some cases be more informative than using solely the cutting efficiency.

## DISCUSSION

In recent years, CRISPR has been used to edit genes and specifically to silence them. The prevalent paradigm is to use computational tools which estimate the likelihood of a mutation following the use of CRISPR and choose a site for gene silencing by taking the site with the highest mutation likelihood. However, when using CRISPR we are usually interested in affecting the expression of the target gene without affecting any other gene’s expression. Thus, the current approaches for designing gRNA do not optimize the right objective.

Therefore, we created EXPosition, a tool which predicts the most likely mutations following CRISPR use and their effect on transcription, splicing and translation initiation. In addition, EXPosition can analyze manually inserted mutations, regardless of their origin, and assess their effects on gene expression. This versatility allows users to assess the effects of mutations that may not have been generated via CRISPR or other specific methods, expanding the tool’s applicability to a wider range of scenarios. Since our tool is composed of various algorithms which predict various aspects of gene expression that were validated and compared to measurements of gene expression, we expect predictions used in our tool to be relevant and correspond with actual expression measurements. We validated our tool using experimental data on thousands of sites and showed our tool can differentiate between sites that will most likely silence a gene and sites that are least likely to do so. We also gave additional information about our outputs by providing the fraction of pathogenic mutations from the ClinVar dataset out of all pathogenic and benign mutations that received certain values of EXPosition score.

We hope that the user-friendly GUI will encourage people to use our tool in their scientific endeavors. We believe that since EXPosition is modular, it will be possible to update each part of the tool with newer and better models, including models that are specific for different cell types and/or Cas proteins.

The study reported here clearly demonstrates two important gaps in the field of CRISPR research: 1) We should carefully design better objective functions to correctly evaluate gRNAs; and 2) We should conduct more experiments that include the target gene in their endogenous genomic context while measuring the effect on gene expression in addition to cutting efficiency. Studies including these data will facilitate better understanding of a given gRNA’s phenotypical effect on its target site, rather than only its genotypical effect.

## DATA AVAILABILITY

Annotations were downloaded from Ensembl. The data we analyzed by Doench et al. can be found at https://doi.org/10.1038/nbt.3437. The ClinVar dataset can be found at https://www.ncbi.nlm.nih.gov/clinvar/

## FUNDING

S.C, S.B and N.L are supported by a fellowship from the Edmond J. Safra Center for Bioinformatics at Tel Aviv University. The study was also supported by the CRISPR-IL consortium.

## ACKNOWLEDGEMENTS

The authors would like to thank Nadav Kra-Oz for contributing to the GUI.

## Notes

### Competing Interest Statement

The authors have declared no competing interest.

## References

1. Pickar-Oliver, A. and Gersbach, C.A. (2019) The next generation of CRISPR–Cas technologies and applications. Nat Rev Mol Cell Biol, 20, 490–507.

2. Uehara, H., Zhang, X., Pereira, F., Narendran, S., Choi, S., Bhuvanagiri, S., Liu, J., Ravi Kumar, S., Bohner, A., Carroll, L., et al. (2021) Start codon disruption with CRISPR/Cas9 prevents murine Fuchs’ endothelial corneal dystrophy. Elife, 10, e55637.

3. Si, X., Zhang, H., Wang, Y., Chen, K. and Gao, C. (2020) Manipulating gene translation in plants by CRISPR–Cas9-mediated genome editing of upstream open reading frames. Nat Protoc, 15, 338–363.

4. Whitworth, K.M., Benne, J.A., Spate, L.D., Murphy, S.L., Samuel, M.S., Murphy, C.N., Richt, J.A., Walters, E., Prather, R.S. and Wells, K.D. (2017) Zygote injection of CRISPR/Cas9 RNA successfully modifies the target gene without delaying blastocyst development or altering the sex ratio in pigs. Transgenic Res, 26, 97–107.

5. Chari, R., Yeo, N.C., Chavez, A. and Church, G.M. (2017) sgRNA Scorer 2.0: a species-independent model to predict CRISPR/Cas9 activity. ACS Synth Biol, 6, 902–904.

6. Listgarten, J., Weinstein, M., Kleinstiver, B.P., Sousa, A.A., Joung, J.K., Crawford, J., Gao, K., Hoang, L., Elibol, M. and Doench, J.G. (2018) Prediction of off-target activities for the end-to-end design of CRISPR guide RNAs. Nat Biomed Eng, 2, 38–47.

7. Lei, Y., Lu, L., Liu, H.-Y., Li, S., Xing, F. and Chen, L.-L. (2014) CRISPR-P: a web tool for synthetic single-guide RNA design of CRISPR-system in plants. Mol Plant, 7, 1494–1496.

8. Liu, H., Wei, Z., Dominguez, A., Li, Y., Wang, X. and Qi, L.S. (2015) CRISPR-ERA: a comprehensive design tool for CRISPR-mediated gene editing, repression and activation. Bioinformatics, 31, 3676–3678.

9. Concordet, J.-P. and Haeussler, M. (2018) CRISPOR: intuitive guide selection for CRISPR/Cas9 genome editing experiments and screens. Nucleic Acids Res, 46, W242–W245.

10. Peng, D. and Tarleton, R. (2015) EuPaGDT: a web tool tailored to design CRISPR guide RNAs for eukaryotic pathogens. Microb Genom, 1.

11. Montague, T.G., Cruz, J.M., Gagnon, J.A., Church, G.M. and Valen, E. (2014) CHOPCHOP: a CRISPR/Cas9 and TALEN web tool for genome editing. Nucleic Acids Res, 42, W401–W407.

12. Shen, M.W., Arbab, M., Hsu, J.Y., Worstell, D., Culbertson, S.J., Krabbe, O., Cassa, C.A., Liu, D.R., Gifford, D.K. and Sherwood, R.I. (2018) Predictable and precise template-free CRISPR editing of pathogenic variants. Nature, 563, 646–651.

13. Stemmer, M., Thumberger, T., del Sol Keyer, M., Wittbrodt, J. and Mateo, J.L. (2015) CCTop: an intuitive, flexible and reliable CRISPR/Cas9 target prediction tool. PLoS One, 10, e0124633.

14. Allen, F., Crepaldi, L., Alsinet, C., Strong, A.J., Kleshchevnikov, V., de Angeli, P., Páleníková, P., Khodak, A., Kiselev, V. and Kosicki, M. (2019) Predicting the mutations generated by repair of Cas9-induced double-strand breaks. Nat Biotechnol, 37, 64–72.

15. Labuhn, M., Adams, F.F., Ng, M., Knoess, S., Schambach, A., Charpentier, E.M., Schwarzer, A., Mateo, J.L., Klusmann, J.-H. and Heckl, D. (2018) Refined sgRNA efficacy prediction improves large-and small-scale CRISPR–Cas9 applications. Nucleic Acids Res, 46, 1375–1385.

16. Moreno-Mateos, M.A., Vejnar, C.E., Beaudoin, J.-D., Fernandez, J.P., Mis, E.K., Khokha, M.K. and Giraldez, A.J. (2015) CRISPRscan: designing highly efficient sgRNAs for CRISPR-Cas9 targeting in vivo. Nat Methods, 12, 982–988.

17. Listgarten, J., Weinstein, M., Kleinstiver, B.P., Sousa, A.A., Joung, J.K., Crawford, J., Gao, K., Hoang, L., Elibol, M. and Doench, J.G. (2018) Prediction of off-target activities for the end-to-end design of CRISPR guide RNAs. Nat Biomed Eng, 2, 38–47.

18. Pulido-Quetglas, C., Aparicio-Prat, E., Arnan, C., Polidori, T., Hermoso, T., Palumbo, E., Ponomarenko, J., Guigo, R. and Johnson, R. (2017) Scalable design of paired CRISPR guide RNAs for genomic deletion. PLoS Comput Biol, 13, e1005341.

19. Heigwer, F., Kerr, G. and Boutros, M. (2014) E-CRISP: fast CRISPR target site identification. Nat Methods, 11, 122–123.

20. Li, V.R., Zhang, Z. and Troyanskaya, O.G. (2021) CROTON: an automated and variant-aware deep learning framework for predicting CRISPR/Cas9 editing outcomes. Bioinformatics, 37, i342– i348.

21. Molla, K.A. and Yang, Y. (2020) Predicting CRISPR/Cas9-Induced Mutations for Precise Genome Editing. Trends Biotechnol, 38, 136–141.

22. Chuai, G., Ma, H., Yan, J., Chen, M., Hong, N., Xue, D., Zhou, C., Zhu, C., Chen, K., Duan, B., et al. (2018) DeepCRISPR: optimized CRISPR guide RNA design by deep learning. Genome Biol, 19, 80.

23. Leenay, R.T., Aghazadeh, A., Hiatt, J., Tse, D., Roth, T.L., Apathy, R., Shifrut, E., Hultquist, J.F., Krogan, N., Wu, Z., et al. (2019) Large dataset enables prediction of repair after CRISPR–Cas9 editing in primary T cells. Nat Biotechnol, 37, 1034–1037.

24. Ben-Yehezkel, T., Zur, H., Marx, T., Shapiro, E. and Tuller, T. (2013) Mapping the translation initiation landscape of an S. cerevisiae gene using fluorescent proteins. Genomics, 102, 419– 429.

25. Agarwal, V. and Shendure, J. (2020) Predicting mRNA Abundance Directly from Genomic Sequence Using Deep Convolutional Neural Networks. Cell Rep, 31, 107663.

26. Jaganathan, K., Kyriazopoulou Panagiotopoulou, S., McRae, J.F., Darbandi, S.F., Knowles, D., Li, Y.I., Kosmicki, J.A., Arbelaez, J., Cui, W., Schwartz, G.B., et al. (2019) Predicting Splicing from Primary Sequence with Deep Learning. Cell, 176, 535-548.e24.

27. Konstantakos, V., Nentidis, A., Krithara, A. and Paliouras, G. (2022) CRISPRedict: a CRISPR-Cas9 web tool for interpretable efficiency predictions. Nucleic Acids Res, 50, W191–W198.

28. Chen, W., McKenna, A., Schreiber, J., Haeussler, M., Yin, Y., Agarwal, V., Noble, W.S. and Shendure, J. (2019) Massively parallel profiling and predictive modeling of the outcomes of CRISPR/Cas9- mediated double-strand break repair. Nucleic Acids Res, 47, 7989–8003.

29. Nicolas Lynn and Tamir Tuller (2022) Detecting and understanding meaningful cancerous mutations based on computational models of mRNA splicing. Under Review.

30. Zhang, S., Hu, H., Jiang, T., Zhang, L. and Zeng, J. (2017) TITER: predicting translation initiation sites by deep learning. Bioinformatics, 33, i234–i242.

31. Cunningham, F., Allen, J.E., Allen, J., Alvarez-Jarreta, J., Amode, M.R., Armean, I.M., Austine-Orimoloye, O., Azov, A.G., Barnes, I., Bennett, R., et al. (2022) Ensembl 2022. Nucleic Acids Res, 50, D988–D995.

32. Landrum, M.J., Lee, J.M., Benson, M., Brown, G.R., Chao, C., Chitipiralla, S., Gu, B., Hart, J., Hoffman, D. and Jang, W. (2018) ClinVar: improving access to variant interpretations and supporting evidence. Nucleic Acids Res, 46, D1062–D1067.

33. Doench, J.G., Fusi, N., Sullender, M., Hegde, M., Vaimberg, E.W., Donovan, K.F., Smith, I., Tothova, Z., Wilen, C., Orchard, R., et al. (2016) Optimized sgRNA design to maximize activity and minimize off-target effects of CRISPR-Cas9. Nat Biotechnol, 34, 184–191.

34. Tuller, T. and Zur, H. (2015) Multiple roles of the coding sequence 5′ end in gene expression regulation. Nucleic Acids Res, 43, 13–28.

35. Ben-Yehezkel, T., Zur, H., Marx, T., Shapiro, E. and Tuller, T. (2013) Mapping the translation initiation landscape of an S. cerevisiae gene using fluorescent proteins. Genomics, 102, 419– 429.

36. Lang, G.I., Murray, A.W. and Botstein, D. (2009) The cost of gene expression underlies a fitness trade-off in yeast. Proceedings of the National Academy of Sciences, 106, 5755–5760.

37. Keren, L., Hausser, J., Lotan-Pompan, M., Vainberg Slutskin, I., Alisar, H., Kaminski, S., Weinberger, A., Alon, U., Milo, R. and Segal, E. (2016) Massively Parallel Interrogation of the Effects of Gene Expression Levels on Fitness. Cell, 166, 1282-1294.e18.

