## Supplementary information for "A tool for CRISPR-Cas9 gRNA evaluation based on computational models of gene expression"

### Supplementary methods

#### 1. Xpresso

Xpresso (1) was trained and tested on cap analysis gene expression (CAGE) data of all genes (~18k). The network consists of ~112k parameters and its architecture is made up of two sequential convolutional and max-pooling layers, after which come two fully connected layers before connecting to the output neuron (Fig. S1). Xpresso was had an $r^2$ of 0.59 when comparing predicted and measured mRNA levels in Human.

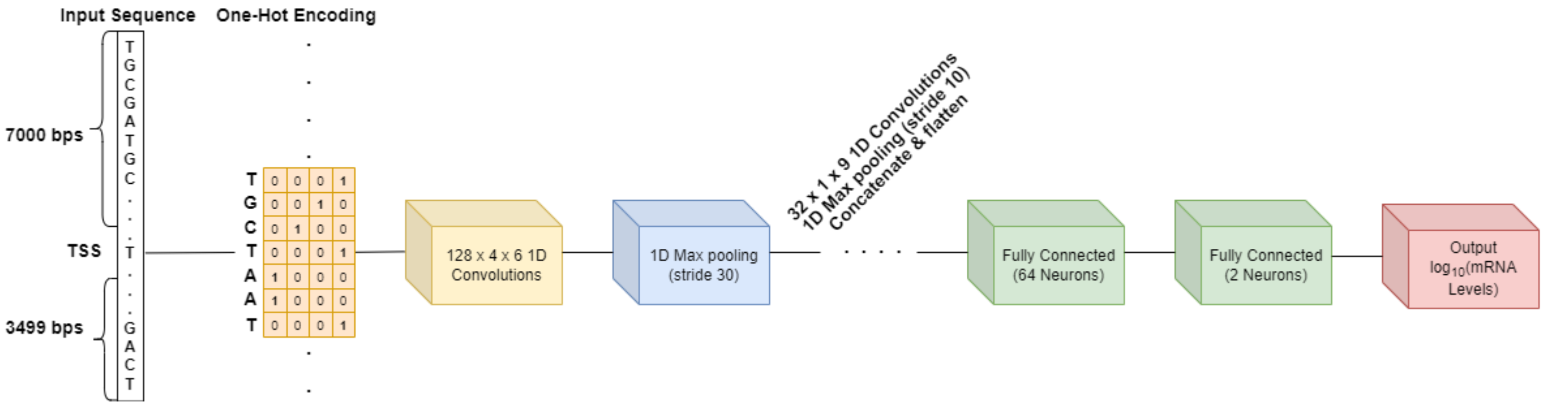


***Fig S1: Schematic of Xpresso.*** *The neural network used to predict mRNA levels from a sequence surrounding the Transcription Start Site (TSS).*

### 2. SpliceAI

The input to SpliceAI (2) is a sequence of at least 10,001nt where each position of interest has at least 10,000nt of context (5000 from each side), and it will output a sequence of acceptor and donor probabilities for all positions of interest i.e. any nucleotide in the middle of the sequence that has 5000nt of context from each side. The network was trained on GENCODE-annotated pre-mRNA transcript sequences (3) on a subset of the human chromosomes to train the parameters of the neural network, and transcripts on the remaining chromosomes (paralogs excluded) to test the network's predictions. The architecture consists of 32 dilated convolutional layers (Fig. S2). For pre-mRNA transcripts in the test dataset, the network predicts spliced positions with 95% top-k accuracy, which is the fraction of correctly predicted splice sites at the threshold where the number of predicted sites is equal to the actual number of splice sites present in the test dataset (4, 5) and has a PR-AUC of 0.98. In addition, the network predicted known splice junctions in long noncoding RNAs (lincRNAs) with 84% top-k accuracy.

For a mutated gene, we input a ~20k nucleotide-long sequence surrounding the start position of the mutation to SpliceAI (depends on the mutation) so that every position that can potentially be affected by the mutation is analyzed (every position up to 5000nt from a mutation is affected, and it will need 5000nt from each side to be analyzed). Therefore, passing this sequence to SpliceAI will return two vectors of ~10k probabilities – one for acceptor sites and another for donor sites.

Note that if the sequence's length is greater than the length of the pre-mRNA it's working on, we pad the input sequence with N's on the sides to make it so every position to be analyzed has 10,000nt of context.

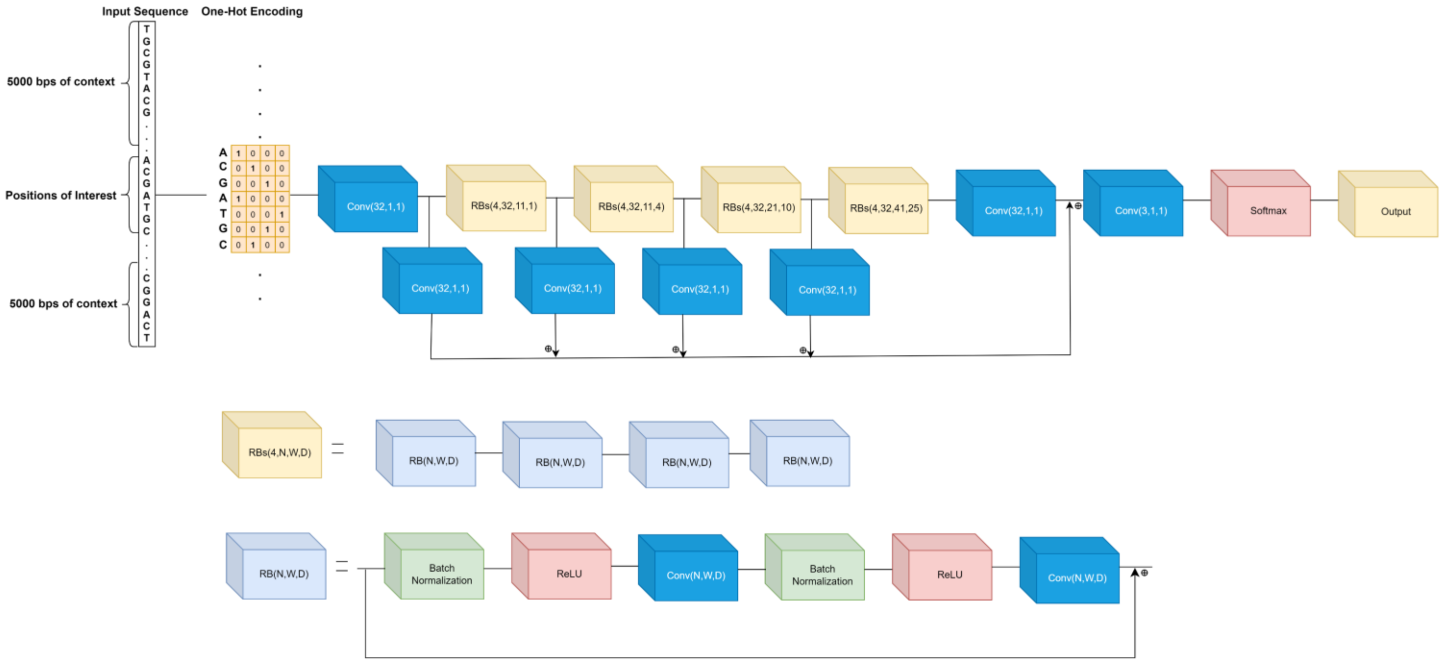


***Fig S2: Architecture of SpliceAI.*** *Each residual block (RB) has three hyper-parameters N, W, and D, where N denotes the number of convolutional kernels, W denotes the window size, and D denotes the dilation rate of each convolutional kernel. The hyper-parameters were chosen such that the number of nucleotides around a position to be analyzed is 10000.*

#### 3. Modeling variant mRNAs of mutated genes

We denote each analyzed position as a missed splice site if it was annotated as a donor/acceptor and its probability of being such a site, as predicted by SpliceAI, has decreased by more than 50%. Conversely, if a position's probability of being a splice site increased by more than 50%, we denote it as a discovered splice site (Fig. S3A).

Following the predicted change in splice sites, we add the discovered splice sites to the annotated sites set and remove the missed splice sites. We then consider all possible isoforms created by this set when sequentially joining splicing sites from the 5' end of the transcript to its 3' end, joining two sites only if one of them is a donor and the other is an acceptor (Fig. S3B). In addition, the isoforms must begin with a donor and end with an acceptor.

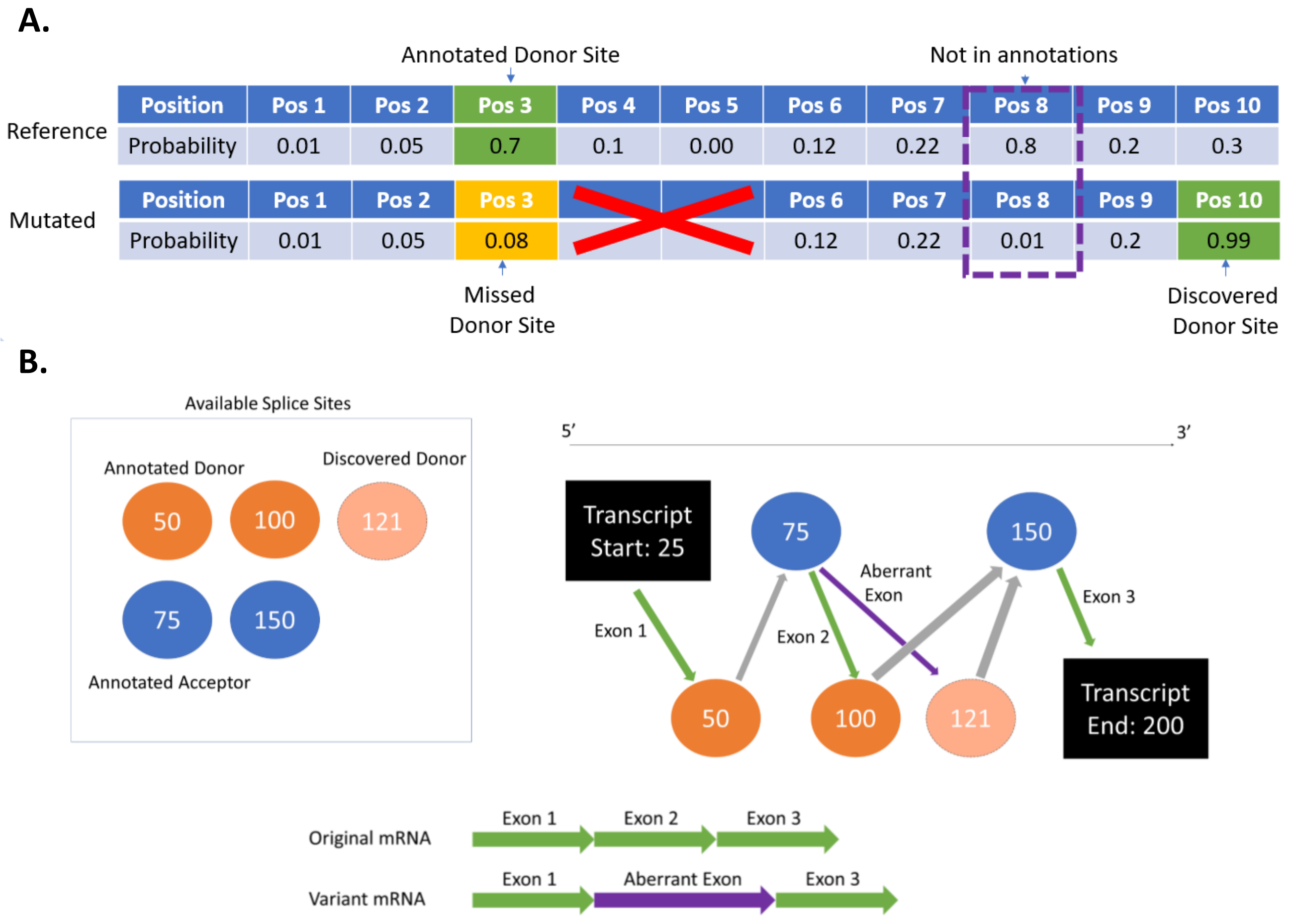


***Fig S3: How isoforms are constructed following mis-splicing. A.*** *An example of how aberrant splicing events are found. Green/yellow cells indicate found/missed splicing sites, respectively. Positions 4-5 were deleted by CRISPR; position 3 is a missed splicing site, since it was annotated as a splicing site and its predicted probability of being a donor site decreased by more than 0.5. Position 10 is a discovered splicing site, since it did not appear in the annotations and its probability of being a donor site increased by more than 0.5. If a position received a high probability in the reference sequence but is not recognized as a splice site in the annotations, it is ignored when looking for missing splice sites.* ***B.*** *An example of constructing different isoforms of the same mRNA. This transcript originally had*

*3 exons, with 2 acceptor and 2 donor sites ("Original mRNA"). Due to a mutation, a discovered donor site was added as an option. Since we cannot know which donor is more likely, we construct an additional isoform, with the discovered splice site ("Variant mRNA").*

Another criterion considered when composing variant mRNAs is that the discovered exons must not exceed 2000nt since exons are usually less than 200nt in length (~80% of annotated transcripts) and are rarely more than 2000nt (less than 1%). Moreover, such long exons they are biologically unlikely and possibly decomposed by the cell (6, 7). With the variant mRNA constructed, the Splicing sub-model checks if the start and stop codons have been mutated in a way that will change their function. If the Translation Initiation sub-model is selected by the user, then the alternative start codon will be the one it outputs. Otherwise, the start codon is the one in the annotations (and if it's deleted then the protein is deemed to have been erased). In case a stop codon is erased, the stop codon that will be chosen will be the first one encountered in the reading frame.

### 4. **Oncosplice**

Oncosplice predicts the disruption caused to a gene following a change in splicing (8). Its first step is using Rate4Site (9), using as input multiple sequence alignments (MSAs) downloaded from the UCSC website (10) which compare the human proteome to the homologous proteomes of 99 vertebrates. Rate4Site scores were calculated for each position in the proteins included in the MSA, covering approximately 90% of the human proteome.

The process of calculating a gene's splicing score is described here and illustrated in Fig. S4; Oncosplice begins by calculating each isoform's score, and based on these acquire transcript- and gene-based scores.

First, for each position $i$ in every analyzed isoform, the normalized conservation score $s_i$ is defined as:

$$(1)\ s_i = e^{-s_i'}$$

Where $s_i'$ is the Rate4Site score for position $i$. If conservation values are not available for a given protein, each amino acid residue is given a score of $s_i = 1$. Then a sliding window is used to calculate average conservation scores along the protein; the window's length was chosen to be 76, which is the mean domain length of all human proteins (calculated using Interpro (11)). For each window, Oncosplice calculates $c = \frac{C}{C_{max}}$, where $C$ is the average $s_i$ score of the window, and $C_{max}$ is the maximum $C$ of all such windows in the proteome. Oncosplice goes over all indels in the window, relative to the original transcript, and defines each indel's disruption score as $s_{indel} = \max\left(1, \frac{l}{w}\right) \cdot c$, where $l$ is the indel's length and $w$ is the window's length (as described above, $w = 76$). The window's score $s_{window}$ would then be the sum of all such $s_{indel}$, and the transcript's score $s_{transcript}$ is the maximal window score. The final score for an isoform is $\min\left(\frac{\max s_{transcript}}{c}, 1\right)$ , where $c$ is the maximum score registered from the ClinVar analysis. If a transcript of interest is provided, we output this score relating only to that transcript.

Having obtained disruption scores for all isoforms, the corresponding transcript's splicing score was defined as the average disruption score over all its isoforms (assuming isoforms have equal probabilities of appearing); and finally, the gene's splicing score is the maximal score out of all its transcripts (i.e. $s'_{ij}$ from Eq. 2 in the main paper). This score ranges from 0 to 1, with a higher score denoting higher disruption to the mutated protein's function.

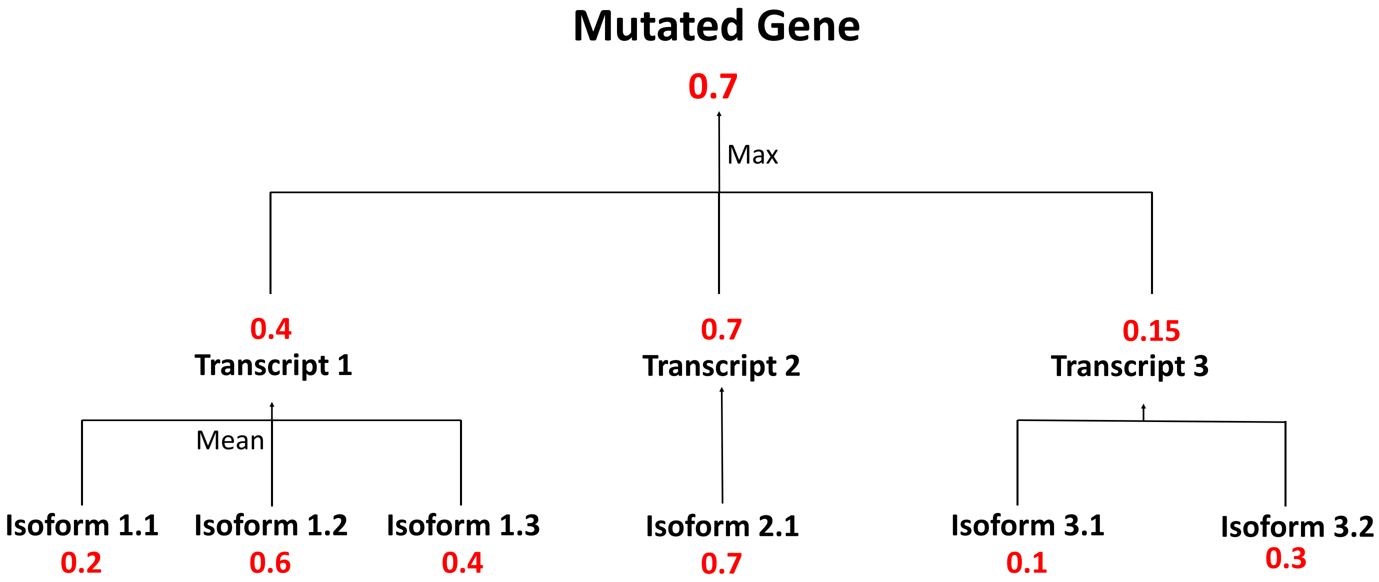


***Fig S4: An example of how the splicing score is calculated for a gene.*** *In this example, following the mutation there are 3 isoforms in Transcript 1, one isoform in Transcript 2, and 2 isoforms in Transcript 3 of the gene. The numbers indicate the splicing score. The score for each transcript is the mean of the scores of the isoforms. The score for the gene is maximum of the scores from the transcripts. Note that this logic applies for all sub-models.*

### 5. TITER

The architecture of the network consists of convolutional layers followed by max pooling, then a recurrent layer of LSTM followed by a logistic layer (Fig. S5). It was trained and tested on QTI-seq data from a HEK293 cell line (12) and the annotated TISs from Ensembl v84 (13) (together denoted by Gao15 from (12)).

When tested on AUG and NUG codon (where N can be any nucleotide) TITER on a dataset constructed from QTI-seq data from (12) which had ~770 positive and ~9900 negative samples, TITER yielded areas under the receiver-operating characteristic and the precision recall curves scores of 89.1 and 61.8% respectively. It's important to note the reason the testing was done only on these codons is that in the QTI-seq data, the authors noticed that these 5 codons made up more than 75% of the cases,

The score for a codon is the product of two terms:

$$(1) \quad TISScore(s) = CodonScore(s) \times ContextScore(s)$$

Where $s$ is the sequence profile of a codon site of interest, $ContextScore(s)$ is the output from the neural network and $CodonScore(s)$ is a term calculated independently to account for the prevalence of certain codons in the human genome (for more details, please see (14))

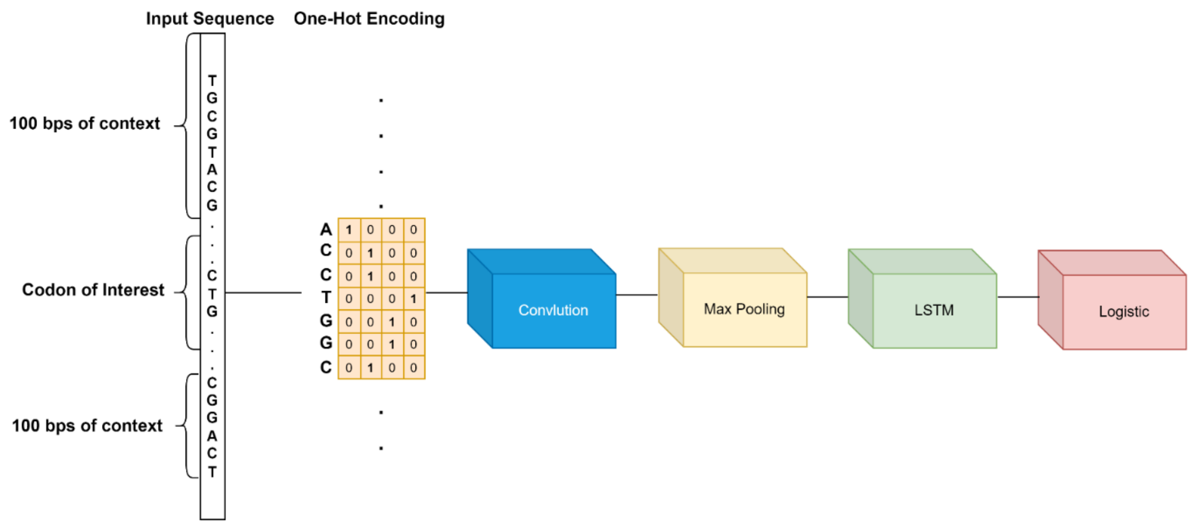


***Fig S5: Schematic illustration of the architecture of TITER.*** *After one-hot encoding the 203-long nucleotide sequence centered around the codon in question, the sequence is passed through several convolution operators, then several pooling operators and then through recurrent layers (LSTM). Finally, the outputs from all the LSTMs are concatenated and fed into a logistic regression layer which outputs a probability of the codon in question to be a TIS.*

### 6. Finding a suitable start codon

For each isoform, we first conduct local alignment (using Biopython) to find the original start codon's location in the mutated mRNA sequence, using a 50nt window centered around the start codon. If the alignment failed (i.e. the score of the best alignment is 0 or less), we say that no start codon was found. Assuming the alignment succeeded, we analyze every possible start codon that is in the same frame as the original mRNA and inside a 40nt window centered around the original start codon. The translation initiation score is calculated using TITER (14), a tool which combines a deep learning algorithm and prior knowledge of preferred TIS codon composition to predict Translation Initiation Sites (TISs); See more details in the "TITER" section. The input to the tool is a 200nt window centered around a potential start codon and the output is a score (≥0) which is indicative of its likelihood to be a TIS. In accordance with the findings described in TITER's paper, the possible start codons we are looking for are NUG (where N is any nucleotide) and ACG.

If any codon has been found and either its TITER score was higher than the original start codon's score, or its rank amongst all TITER scores for the annotated TISs in the human transcriptome was within 5% of the original start codon's rank, then this codon is deemed a suitable start codon; out of all the suitable codons found this way, the new start codon is defined to be the one with the highest TITER score. If we haven't found any suitable start codons, we repeat the process with a 100nt window instead of 40nt; if we still find no suitable codons, we repeat the process with increasing window sizes: 200nt, 400nt and 800nt. If this process still hasn't yielded suitable start codons, the new start codon would be defined as the highest-scoring potential start codon found in the final, 800nt window.

### 7. Chosen thresholds for ClinVar analysis

We list here the 95th-99th percentile values of each EXPosition sub-models' score; see "High-scoring mutations are overrepresented in ClinVar-designated pathogenic mutations" and figure 3 in the main text.

| Percentile | Transcription Score | Splicing Score | Translation Initiation Score |
|---|---|---|---|
| 0.99 | 0.119 | 6687 | 0.778 |
| 0.98 | 0.076 | 5060 | 0.435 |
| 0.97 | 0.056 | 4096 | 0.099 |
| 0.96 | 0.044 | 3388 | 0.026 |
| 0.95 | 0.035 | 2880 | 0.005 |

***Table S1: Raw scores of sub-models in ClinVar analysis corresponding to the percentiles 95%-99%.***

In table S2, we provide the maximal scores calculated by each sub-model on the ClinVar set, as well as the raw and normalized threshold values used in the final program.

| | Transcription | Splicing | Translation initiation |
|---|---|---|---|
| **Raw threshold** | 0.02 | 21 | 0.099 |
| **Maximal value** | 0.97 | 105,944 | 65.42 |
| **Normalized threshold** | 0.02 | $1.98 \times 10^{-4}$ | 0.001 |

***Table S2: Raw threshold, maximal value achieved and normalized threshold values for each sub-model***

It is also possible for the user to increase the sensitivity of the tool by taking the maximum score (instead of the average) over all mutations predicted\inserted and checking if it passes the threshold. This is done by checking the appropriate box in the tool.

### 8. Averaging of repeats in the Doench et al. experiment

For the A375 data, we first averaged the sgRNA fold change over the repeats, separately for each delivery system subset (lentiCRISPRv2 and lentiGuide); we then set the two subsets to have the same mean by multiplying all measurements in one of them with a constant. Finally, for each target site we took the average sgRNA fold change over the two delivery subsets. For the HT29 data, we simply took the average over all repeats.

### 9. The predictive power of repeats of the Doench et al. experiment

For each cell and delivery method, we trained SVM classifiers whose single feature was the raw data from one of its repeats and the labels for the least/most silencing site were chosen using the raw data from another repeat. For each gene, we took the most and least silencing sites (according to the fold change it incurred) based on one of the repeats and labeled them accordingly; we then used each site's fold change, based on one of the other repeats, as a feature in the classifier. We trained the classifiers on randomly selected 80% (~28.8k) of the sites and tested the trained classifier in each case on the remaining 20% (~7.2k) of the sites. We performed this 100 times, randomly selecting the training/test sets each time.

The mean loss over the 100 repetitions can be seen in Fig. S6:

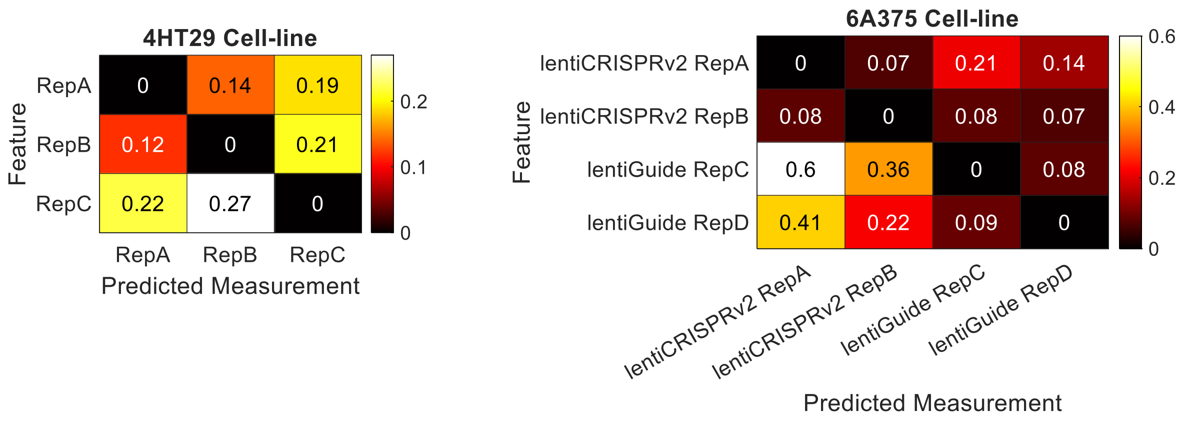


***Fig S6: Heatmaps of the average losses when training on empirical data.*** Mean loss values, over 100 randomized repetitions, for SVM models trained using one experiment repeat (vertical axis) to predict the most\least silencing sites according to another repeat (horizontal axis).

The losses between the repeats in the HT29 cells were larger by up to two-fold compared to the losses between the repeats in the A375 cells, in the same delivery systems. The fact the loss is higher between delivery systems in the A375 cells (in the 4 top-right and 4 bottom-left boxes) further supports our claim for the biological stochasticity of the data.

The mean loss when training on empirical data was 0.43 (Fig. 6C) while the mean loss when training with EXPosition was 0.47. This leads us to believe that the data was noisy and that we gained satisfactory results seeing as they were almost as optimal as the results using empirical data.

### References


1. Agarwal,V. and Shendure,J. (2020) Predicting mRNA Abundance Directly from Genomic Sequence Using Deep Convolutional Neural Networks. *Cell Rep*, **31**, 107663.

2. Jaganathan,K., Kyriazopoulou Panagiotopoulou,S., McRae,J.F., Darbandi,S.F., Knowles,D., Li,Y.I., Kosmicki,J.A., Arbelaez,J., Cui,W., Schwartz,G.B., *et al.* (2019) Predicting Splicing from Primary Sequence with Deep Learning. *Cell*, **176**, 535-548.e24.

3. Harrow,J., Frankish,A., Gonzalez,J.M., Tapanari,E., Diekhans,M., Kokocinski,F., Aken,B.L., Barrell,D., Zadissa,A. and Searle,S. (2012) GENCODE: the reference human genome annotation for The ENCODE Project. *Genome Res*, **22**, 1760–1774.

4. Yeo,G. and Burge,C.B. (2003) Maximum entropy modeling of short sequence motifs with applications to RNA splicing signals. In *Proceedings of the seventh annual international conference on Research in computational molecular biology*.pp. 322–331.

5. Boyd,S., Cortes,C., Mohri,M. and Radovanovic,A. (2012) Accuracy at the top. *Adv Neural Inf Process Syst*, **25**.

6. Singh,P., Saha,U., Paira,S. and Das,B. (2018) Nuclear mRNA Surveillance Mechanisms: Function and Links to Human Disease. *J Mol Biol*, **430**, 1993–2013.

7. Garneau,N.L., Wilusz,J. and Wilusz,C.J. (2007) The highways and byways of mRNA decay. *Nat Rev Mol Cell Biol*, **8**, 113–126.

8. Nicolas Lynn and Tamir Tuller (2022) Detecting and understanding meaningful cancerous mutations based on computational models of mRNA splicing. *Under Review*.

9. Pupko,T., Bell,R.E., Mayrose,I., Glaser,F. and Ben-Tal,N. (2002) Rate4Site: an algorithmic tool for the identification of functional regions in proteins by surface mapping of evolutionary determinants within their homologues. *Bioinformatics*, **18**, S71–S77.

10. Pupko,T., Bell,R.E., Mayrose,I., Glaser,F. and Ben-Tal,N. (2002) Rate4Site: an algorithmic tool for the identification of functional regions in proteins by surface mapping of evolutionary determinants within their homologues. *Bioinformatics*, **18**, S71–S77.

11. Paysan-Lafosse,T., Blum,M., Chuguransky,S., Grego,T., Pinto,B.L., Salazar,G.A., Bileschi,M.L., Bork,P., Bridge,A., Colwell,L., *et al.* (2023) InterPro in 2022. *Nucleic Acids Res*, **51**, D418–D427.

12. Gao,X., Wan,J., Liu,B., Ma,M., Shen,B. and Qian,S.-B. (2015) Quantitative profiling of initiating ribosomes in vivo. *Nat Methods*, **12**, 147–153.

13. Aken,B.L., Ayling,S., Barrell,D., Clarke,L., Curwen,V., Fairley,S., Fernandez Banet,J., Billis,K., García Girón,C. and Hourlier,T. (2016) The Ensembl gene annotation system. *Database*, **2016**.

14. Zhang,S., Hu,H., Jiang,T., Zhang,L. and Zeng,J. (2017) TITER: predicting translation initiation sites by deep learning. *Bioinformatics*, **33**, i234–i242.
